# A Deep Learning-based Method for Drug Molecule Representation and Property Prediction

**DOI:** 10.1101/2025.07.15.665023

**Authors:** Qi Zhang, Xuan Yu, Yuxiao Wei, Zhi-Hui Wang, Dong-Jun Yu

## Abstract

Accurately and robustly representing drug molecule features, prediction of drug-target biomacromolecule interactions, and determining drug molecule physicochemical properties are crucial in drug development. However, due to issues such as insufficient generalization ability of single-modal representation, lack of multi-task prediction frameworks, and weak adaptability in cold-start scenarios, these tasks remain challenging. Here, we introduce DrugDL, a framework designed for drug molecule representation and the prediction of multiple downstream tasks, including drug-target interactions, binding affinities, binding sites, physicochemical properties, toxicity, and drug-drug interactions. DrugDL achieves joint representation learning of the drug chemical space and the target protein biological space and analyzes the multi-scale interaction mechanisms between drug molecules and target proteins by introducing cross-modal contrastive learning and single-modal feature enhancement algorithms. It employs a multi-task prediction framework to predict multiple properties of drug molecules. In practical applications, DrugDL outperforms state-of-the-art methods, especially in cold-start tasks. It’s successfully applied to high-throughput screening, identifying inhibitors for SARS-CoV-2 and metabolic enzymes, and aids in predicting cancer-targeted drugs. Validations for EGFR and ALK targets confirm its efficiency as a precise drug discovery tool. Leveraging accurate molecular representation and multi-property prediction, DrugDL provides full-chain technical support for drug development, significantly accelerating the drug discovery process.

## Introduction

With the rapid advancement of biomedicine, drug R&D is pivotal for enhancing human health and driving industrial innovation^1^. At its core, it involves decoding the interactions between drug molecules and biomacromolecules like proteins and DNA, as well as understanding the physical and chemical properties of drug molecules^2,3^. These aspects, including interaction patterns, target binding modes, and toxicity, determine key drug parameters, forming the basis for screening and optimization. While experimental methods are crucial in drug discovery, biochemical experiments for large-scale interaction identification and property determination are costly and time-consuming^4,5^. Thus, computational methods have been extensively adopted in drug discovery, achieving notable progress in predicting drug properties, effectively shortening the R&D cycle and cutting costs^6,7^.

The core of computational methods applied to drug property prediction lies in exploring more efficient and reasonable methods for representing the features of drugs (and target biomolecules) and developing better predictors^8,9^. With the rapid development of computer technology and the field of deep learning, revolutionary changes have occurred in molecular feature representation methods^10^. Existing representation methods can be roughly divided into three categories: text-based representation^11,12^, fingerprint-based representation^13^, and graph-based representation**^Error! Reference source not found.^**,^15^. Text-based representation encodes drug molecule structural information as text to capture basic features. There are diverse text-based representation methods for drug molecules, among which the simplified molecular-input line-entry system (SMILES) representation and InChI representation^12^ are the most widely used. A standardized language, SMILES uses short strings to represent chemical structures, offering simplicity, uniqueness, reversibility, and serving as input for most drug property prediction models. Fingerprint-based drug molecule representation converts chemical structure information into a 1D binary vector, where each bit indicates the presence or absence of a specific structural feature. Currently, common molecular fingerprint features include MACCS fingerprints^16^, and ECFP fingerprints^17,18^, etc. Graph-based representation learning of molecules has received increasing attention in recent years and can be roughly divided into two major categories: molecular graphs^19,**Error!**^ **^Reference source not found.^** and molecular images^21^. These representation methods usually rely on specific software or libraries, such as RDkit. They can parse SMILES strings, identify atomic and chemical bond information therein, and construct molecular graphs and generate images accordingly. A molecular graph is an abstract representation method based on nodes and edges, focusing on showing the structural skeleton of drug molecules and the connection relationships between atoms^22^. With the continuous in-depth research on drug molecule substructures and functional groups, drug motif graphs have gradually emerged in many drug property prediction models**^Error! Reference source not found.^**,^23,24^. Different from drug atom graphs, drug motif graphs are a higher-level graph model that focuses on the structural fragments in drug molecules and their interactions. A molecular image is a 2D graphical representation method, usually constructed based on the red, green, and blue color channels, which can show the structural features of molecules in an intuitive and concise manner^25,26^.

When facing complex and diverse drug molecule property prediction tasks, scholars have developed various computational models for different problems. In the prediction of drug-target interactions (DTIs), which is crucial in drug R&D, most methods use the linear and 2D structural information of drugs and targets as input^27^, treat the prediction of DTIs as a binary classification task, and use different deep encoding and decoding modules such as deep neural networks, graph neural networks (GNNs), convolutional neural networks (CNNs), and transformers for prediction^28,29^. Examples of such models include DrugBAN^30^, ZeroBind^31^, PSICHIC^32^, DTIAM^33^, and GraphBAN^34^. With the rapid increase in the number of known target structures and the improvement in the accuracy of AlphaFold-like protein 3D structure prediction methods, a feasible approach has been provided for effectively modeling 3D target structures. Recently, many deep learning models have integrated 3D structure information to improve the prediction of DTIs^35,36^. To further predict the assumed strength of interactions, various regression-based models have been proposed to infer the binding affinities between drugs and targets^37,38^. Binding affinity reflects the degree of binding between drug and specific target and can be quantified by indicators such as the inhibition constant *K_i_*, dissociation constant *K_d_*, and half-maximal inhibitory concentration *IC*_50_. Models for predicting drug-target binding affinity (DTA) and DTI usually adopt similar feature representation and extraction methods, and only modify the final output mapping results and objective functions in the predictors. Examples of such models include KDBNet^39^ and MMD-DTA^40^. Although these methods can successfully predict more detailed affinity strengths compared to DTI-type tasks, their ability to reveal more precise specific binding sites is still very limited. Therefore, many models introduce the attention mechanism to assign weights to features representing different sequence segments according to their importance levels, in order to display specific binding sites. For example, Li et al. developed a method called MONN^41^, which uses non-covalent interactions as additional supervision information to guide the model to capture key binding sites. Hua et al. developed a multi-functional model called MFR-DTA^42^, which uses the Mix-Decoder module and fully connected layers to treat the prediction of drug-target binding regions (DTBRs) as a supervised learning task.

In the prediction of drug molecule physicochemical properties, toxicity, and drug-drug interactions (DDIs), which are safety-related prediction tasks, different from DTI and DTA tasks that involve complex interactions between multi-modal biological entities, most models for predicting these tasks only focus on and use molecular structures, and pay more attention to the action regions and key functional groups of drug molecules^43,44^. These methods have two ways of revealing substructures in drug molecules: explicit and implicit. Explicit models decompose drug molecules according to certain settings and then use a tokenization process similar to that in natural language processing for representation^45^. In contrast, implicit models usually use molecular graphs as input and adopt a combination of GNN and attention mechanisms to automatically extract substructures from each molecular graph^46^. For example, the MolCLR^47^ model proposed by Wang et al. uses GNN, conducts self-supervised learning with a large amount of unlabeled data, and performs molecular property predictions in tasks such as predicting the human β-secretase inhibitors (BACE), blood-brain barrier permeability (BBBP), and drug molecule toxicity, while also identifying important substructures. Zeng et al. used the image representation of drug molecules to detect various physicochemical properties of drug molecules and split the molecular structure by identifying clusters using a multi-granularity chemical cluster classification method^21^. The MeTDDI method proposed by Zhong et al. predicts DDIs through local-global self-attention and co-attention mechanisms and accurately explains the structural mechanism of DDI^48^.

Although a great deal of effort has been invested in the feature representation of drug molecules and the prediction of various molecular properties related to drug R&D, previous studies still have some limitations. Firstly, most representation methods focus on single-modal drug molecule information, restricting their ability to capture deep-level features, especially those related to other biomolecules^9,49^. This leads to poor generalization in predicting new drug-target bindings, akin to the cold start issue. In addition, the existing best drug molecule property prediction methods are often limited to one-sided predictions. For example, some are only related to the prediction of DTI and DTA, while others can only be used for the prediction of DDIs. There is a lack of a comprehensive method that can cover multiple important steps in the drug R&D process. Moreover, previous methods also rely heavily on large-scale high-quality labeled data and interconnected biomedical entities (diseases, and side effects, etc.)^50,51^, but such data is scarce, especially in early drug discovery. More importantly, previous methods fail to elucidate drug action mechanisms and struggle to translate into real-world drug virtual screening, remaining mainly theoretical with limited practical validation of candidate drugs. Developing advanced deep learning models for multi-property prediction, mechanism exploration, and real-world application thus represents a critical yet challenging task in drug development.

In this study, we developed DrugDL, a cross-modal deep learning architecture for drug molecule representation and multi-property prediction. DrugDL takes the SMILES sequences of drug molecules and the amino acid sequences of target as inputs, and constructs drug molecular graphs and motif graphs based on specific rules to represent molecular features at different granularities. DrugDL introduces cross-modal contrastive learning and single-modal feature enhancement algorithms, utilizing contrastive learning loss to align the features of the drug and target modalities, while employing matrix singular value decomposition (SVD) and Gaussian kernel functions to compute intra-modal data similarity and preserve internal modal relationships. Based on this architecture, DrugDL analyzes the multi-scale interaction mechanisms between drugs and targets, achieving joint representation learning of drug chemical space and target biological space. Additionally, it adopts a multi-task prediction framework to test and evaluate various downstream tasks, including DTI, DTA, DTBR, drug physicochemical property, toxicity, and DDI prediction. In the comprehensive comparison tests of various tasks and multiple scenarios (warm start, cold start), DrugDL outperforms state-of-the-art methods in all tasks. In addition, we successfully identified effective inhibitors against SARS-CoV-2 and metabolic enzymes from a high-throughput molecular library and provided theoretical support for drugs in different clinical trial stages by predicting drugs that interact with cancer targets. Moreover, the virtual screening of candidate drugs for EGFR and ALK targets confirm that DrugDL can serve as an efficient and accurate drug discovery tool to assist drug R&D. All these results indicate that DrugDL can accurately represent drug molecules and effectively predict their multi-property, thus greatly facilitating the drug discovery process.

## Results

### Overview of DrugDL

DrugDL is a cross-modal deep learning architecture. It realizes joint representation learning of the drug chemical space and the target protein biological space by analyzing the multi-scale interaction mechanisms between drug molecules and target proteins (Fig. 1 and Extended Data Fig. 1). The model takes the SMILES of drug and the amino acid sequences of target as input. Through a specific substructure splitting method, drug molecules are represented as atomic-level topological graphs and pharmacophore motif graphs, and the target protein sequences are simultaneously mapped into feature matrices. Subsequently, DrugDL uses multi-channel GNNs and multi-layer CNN blocks to learn the embedding representations of drugs and targets, respectively, reflecting the spatial structure features of drug molecules and the long-range dependence features of target protein sequences. Then, the learned embedding representations are combined. Through cross-modal interaction learning, deep fusion of multi-modal information is achieved, and the heterogeneous features of the two modalities of drugs and targets are aligned into a unified feature space. At the same time, by introducing a single-modal preservation loss, the relationships between the internal features of each modality are strengthened, thereby avoiding the risk of overemphasizing inter-modal alignment and neglecting the complexity of the internal feature distribution of each modality when data resources are relatively limited. The drug molecular features extracted by DrugDL are widely applied to multiple downstream tasks, including the prediction of DTI, DTA, DTBR, drug physicochemical properties, drug toxicities, and DDI. The innovativeness and high efficiency of this model provide powerful support for the drug discovery and development process.

**Fig. 1.**
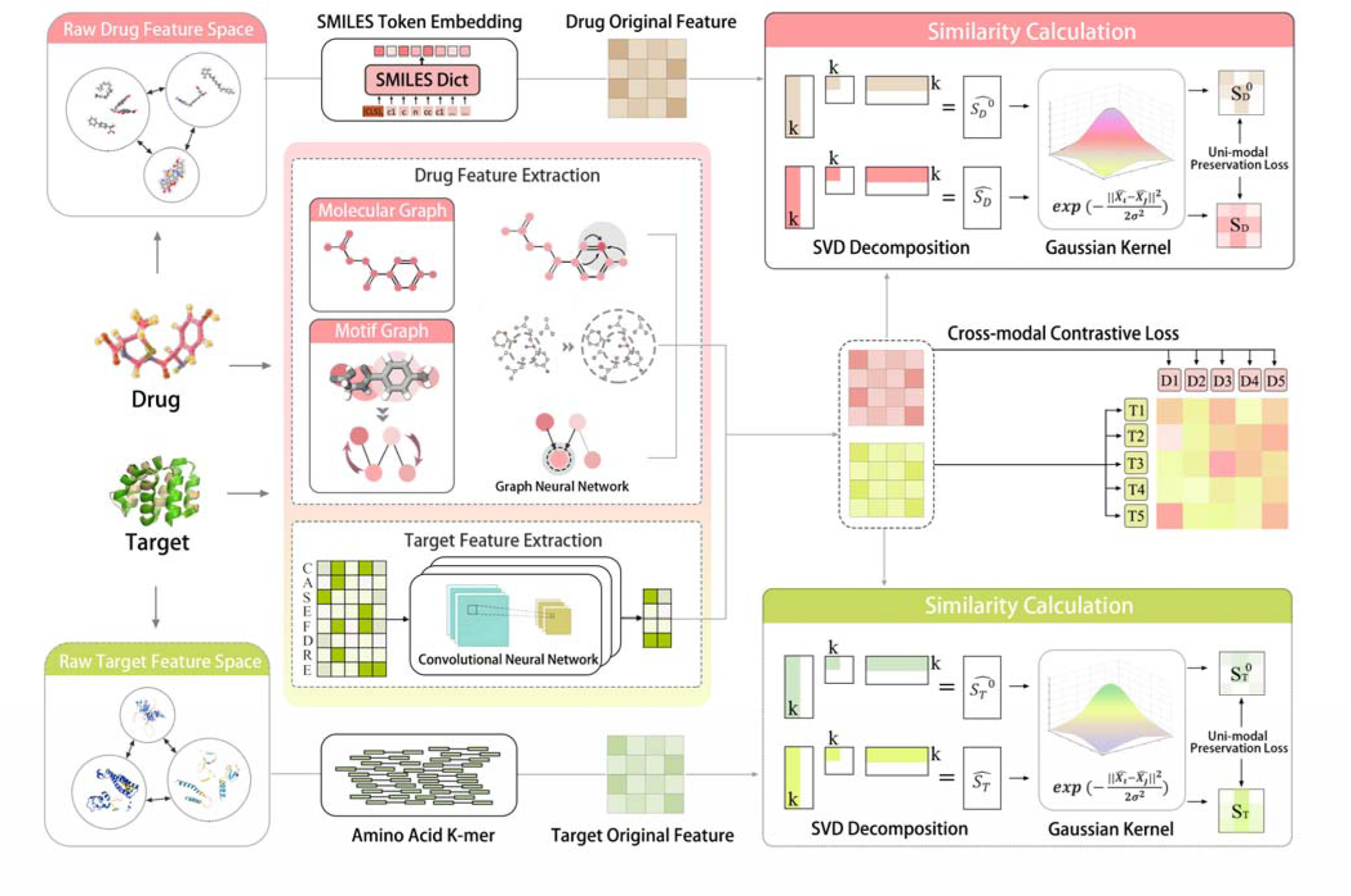
The architecture overview of DrugDL. The framework mainly has two modules. Cross-modal interaction learning module: Converts drug molecules into atomic-level topological and motif graphs, and maps target protein sequences to feature matrices. Uses multi-channel GNNs and multi-layer CNN blocks to learn drug and target embeddings. Integrates embeddings and aligns heterogeneous features into a unified space via cross-modal interaction learning. Single-modal feature enhancement module: Calculates single-modal data similarity with matrix SVD and Gaussian kernel functions. Introduces a preservation loss to strengthen internal feature relations within each modality. Jointly optimizes the two modules for expressive features.

**Extended Data Fig. 1.**
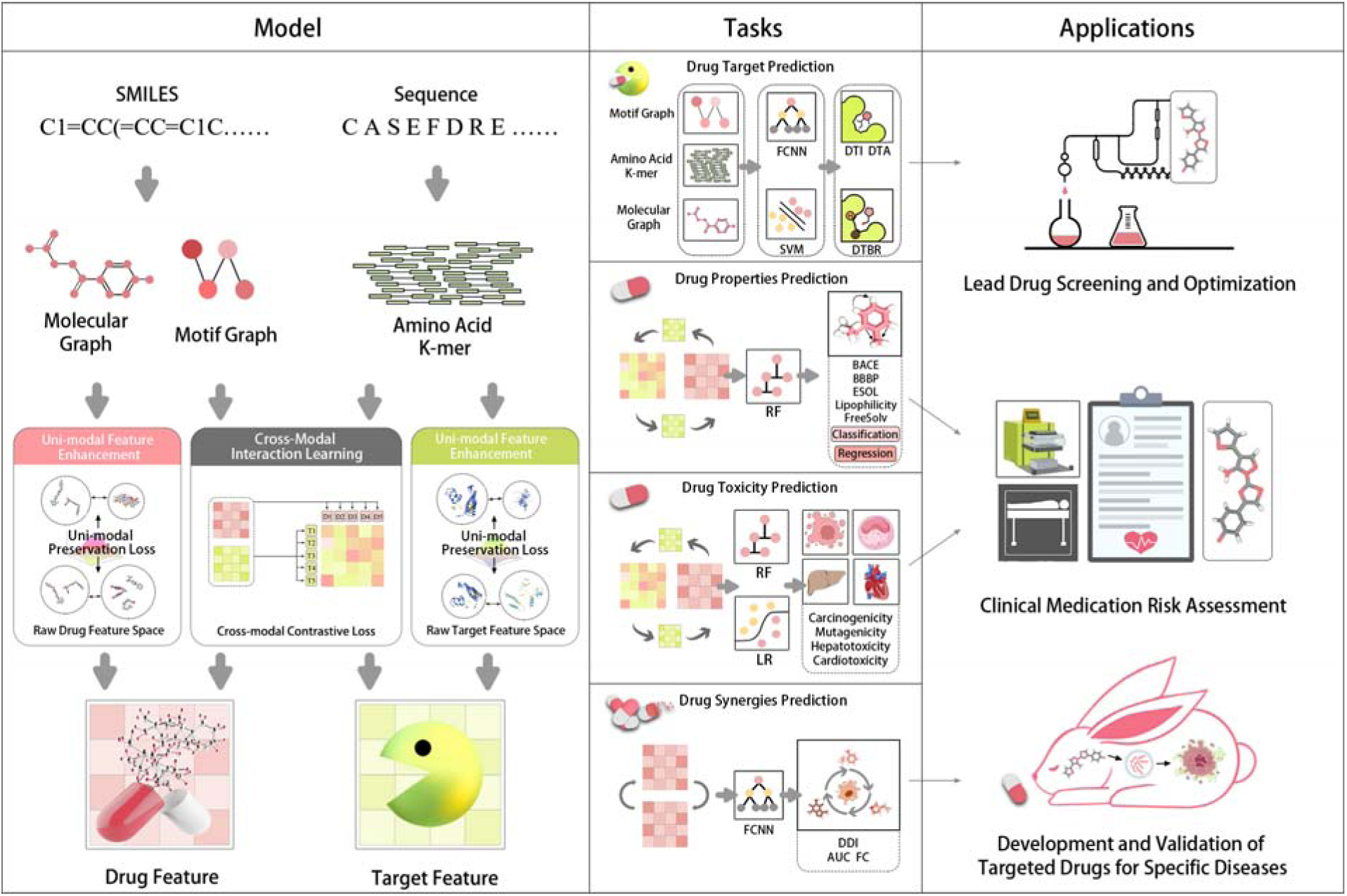
Schematic Diagram of DrugDL Application. The DrugDL model integrates multiple drug and target features and uses cross-modal interaction learning and single-modal feature enhancement techniques for training and testing on benchmark datasets. This model can be applied to various downstream tasks, including the prediction of DTI, DTA, DTBR, and the prediction of molecular properties such as human β-secretase inhibitor (BACE) and blood-brain barrier permeability (BBBP). Moreover, DrugDL can also conduct toxicity predictions such as molecular carcinogenicity, mutagenicity, hepatotoxicity (DILI), and hERG cardiotoxicity, as well as DDI predictions. Through the model’s decision analysis and interpretability analysis, the mechanism of action of drugs can be clarified, suggestions for lead drug optimization can be provided, and the clinical toxicity risks of drug molecules can be evaluated. DrugDL can also perform virtual screening of drug molecules for specific targets and verify the effectiveness and safety of the screened drug molecules through experiments, providing support for the development and verification of therapeutic drugs for specific diseases.

### Performance of DrugDL on the DTI, DTA and DTBR prediction tasks

The DTI prediction task is a classification task, and the goal is to use a model to determine whether a given drug-target pair interacts. In this study, we first combined the features extracted by DrugDL with five common molecular fingerprint features, namely Morgan fingerprints, ECFP fingerprints, PubChem fingerprints, MACCS fingerprints, and Pharmacophore ErG fingerprints. On the benchmark datasets, we utilized the DTI prediction task to reflect the performance capabilities of each feature. As can be seen from Fig. 2a, DrugDL achieved better prediction performance in all evaluation metrics on the DTI benchmark datasets. Compared with other fingerprint features, the DrugDL model demonstrated significant advantages in the Recall and Specificity metrics. We believe that this advantage mainly stems from the imbalance between positive and negative samples in the DTI dataset. In this case, all fingerprint features perform better in the Specificity metric than in the Recall metric. This indicates that when faced with class imbalance, most fingerprint features may mistakenly classify many truly interacting drug-target pairs as negative samples, increasing the risk of missed detections. However, the good performance of the DrugDL model in the Recall and Specificity metrics indicates that in the DTI prediction task, the latent representations extracted by DrugDL can flexibly adapt to changes in the dataset size and effectively address the sample imbalance problem, thus demonstrating strong robustness and reliability. In addition, the t-SNE visualization shown in Fig. 2b is consistent with these results: traditional fingerprints have poor discriminability for positive/negative samples, while DrugDL effectively separates samples into distinct clusters, demonstrating better feature discriminability and robustness.

**Fig. 2.**
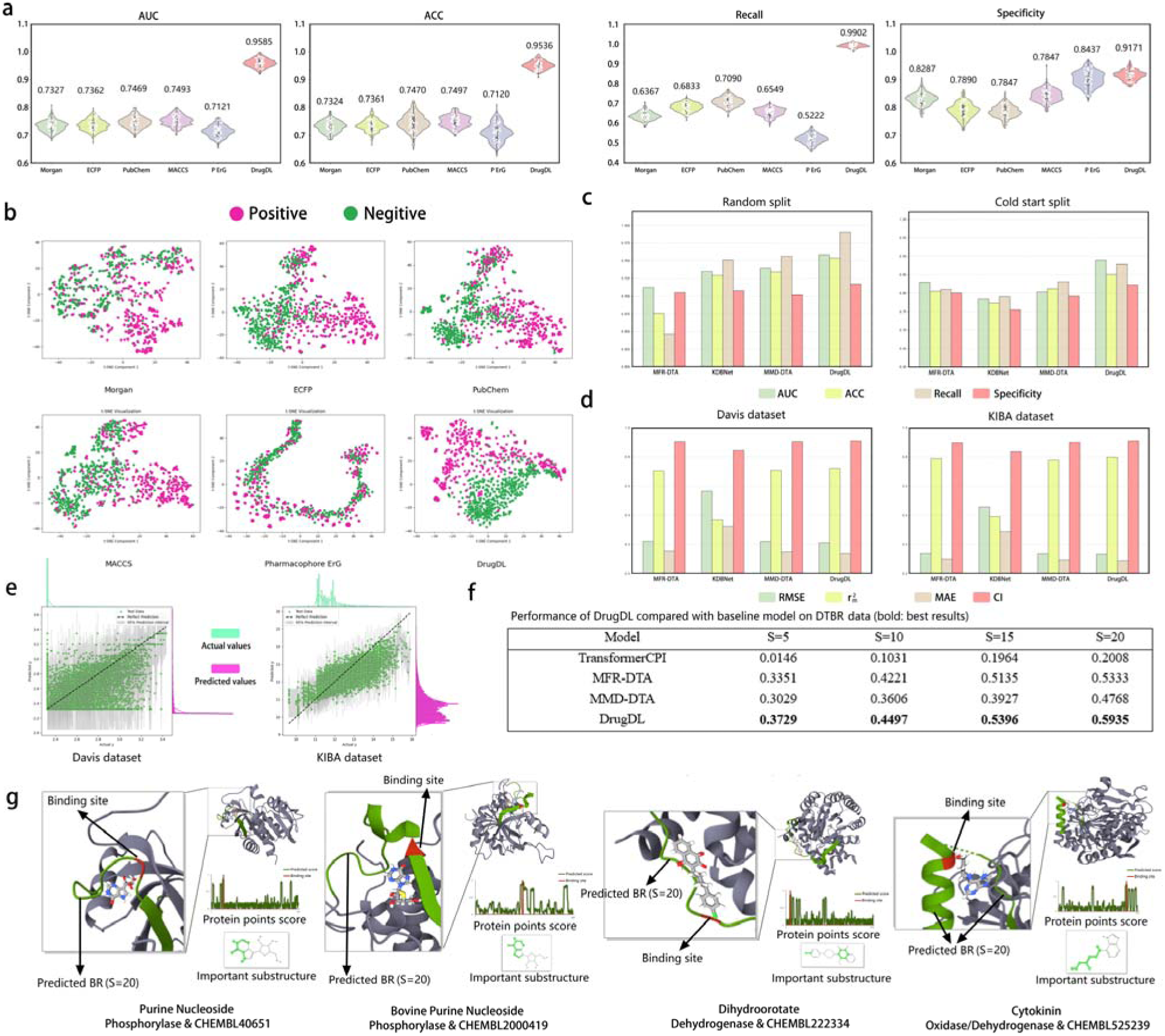
Performance evaluation on the DTI, DTA and DTBR prediction task. **a**, The performance of DrugDL and various molecular fingerprint features was evaluated using AUC, ACC, Recall, and Specificity on the DTI benchmark dataset. **b**, Comparison of t-SNE visualization results of various molecular fingerprint features on the DTI benchmark dataset. **c**, Performance evaluation of DrugDL and baseline models in terms of AUC, ACC, Recall and Specificity on randomly split and cold start split datasets from the DTI benchmark dataset. **d**, Evaluation of the performance of DrugDL and baseline models by RMSE, 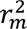, MAE and CI on the Davis and KIBA DTA datasets. **e**, Prediction visualization results of DrugDL on the Davis and KIBA DTA datasets. **f**, Evaluation of the performance of DrugDL and baseline models on the DTBR dataset (sc-PDB). **g**, Prediction visualization results of DrugDL on four complexes in the DTBR (sc-PDB) dataset.

The comparative analysis of DrugDL and the state-of-the-art DTI prediction models (DrugBAN^30^, ZeroBind^31^, and PSICHIC^32^) on randomly split data and cold start split data is shown in Fig. 2c. Obviously, DrugDL significantly outperforms all baseline models not only on randomly split data but also on the more challenging cold start split data, demonstrating excellent performance. In the evaluation of cold start split data, although the performance of all models declined, DrugDL still maintained a relatively leading edge. Its AUC value is 0.8891, much higher than that of DrugBAN (0.8296), ZeroBind (0.7845), and PSICHIC (0.8034). In addition, DrugDL also performed outstandingly in metrics such as ACC, Recall, and Specificity, with values of 0.8506, 0.8787, and 0.8225 respectively, fully demonstrating its strong ability to handle cold start problems. The above results achieved by DrugDL are closely related to its various modules. Therefore, we illustrate in Supplementary Fig. 1 the impact of different feature extraction methods and distinct modules within DrugDL on the model’s prediction outcomes.

In the regression task of DTA prediction, we selected three baseline models specifically designed for the DTA prediction task, MFR-DTA^42^, KDBNet^39^, and MMD-DTA^40^, for an in-depth comparative analysis with DrugDL. Through comprehensive evaluations on the two datasets, Davis and KIBA, we examined the prediction performance of each model in detail. As shown in Fig. 2d, the DrugDL model significantly outperformed the baseline models in all evaluation metrics. Its 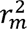 value is much higher than that of other models, fully demonstrating a high degree of correlation between the predicted and actual values. At the same time, the RMSE and MAE values of the DrugDL model also reached the lowest levels among all models, which once again strongly confirms its excellent prediction accuracy on the DTA dataset. In addition, the concordance index CI of DrugDL is relatively high, further enhancing the persuasiveness of the reliability of its prediction results. Fig. 2e intuitively shows the prediction visualization results of DrugDL on the two datasets. The points on its scatter plot are closely distributed around the diagonal, clearly indicating a small deviation between the predicted and actual values. In the KIBA dataset, the label distribution of the data is relatively normal, that is, the actual values of DTAs are more evenly distributed, which provides favorable conditions for the model to learn a more accurate mapping relationship.

In the more refined DTBR prediction task, we selected TransformerCPI^52^, MFR-DTA, and MMD-DTA for performance evaluation and comparative analysis with DrugDL. To objectively assess prediction capabilities, we designed a standard evaluation using the probability that the actual binding site falls within the predicted region as the core metric. We set varying lengths of predicted DTBRs, comprising *S* amino acids (*S =* 5, 10, 15, 20), to evaluate model performance across different segment lengths. The results are shown in Fig. 2f, which details the prediction accuracy of each model under different *S* values. It can be clearly seen from the data in the table that as the value of *S* increases, the accuracy of the predicted binding regions of all methods shows an upward trend. When *S* is equal to 5 and 10, although the prediction accuracy of all baseline models is relatively low, the DrugDL model still demonstrated significant advantages in handling shorter sequence segments. When *S* takes values of 15 and 20, the prediction performance of the DrugDL model was further enhanced, with accuracies reaching 0.5396 and 0.5935 respectively. Compared with other baseline models, the prediction performance of DrugDL is far ahead when *S = 15* and *S = 20*, which once again strongly verifies its excellent effectiveness and high reliability in the DTBRs prediction task. Fig. 2g illustrates the alignment between DrugDL-predicted binding regions (*S = 20*, green) and experimentally verified actual binding sites (red) for four drug-target pairs. For three proteins, predicted binding regions accurately encompass the actual sites. For cytokinin oxidase/dehydrogenase, the predicted site is slightly offset but still within the predicted region. Additionally, we visualized drug molecules based on motif features, with red areas highlighting key substructures in drug-protein binding. These results collectively underscore DrugDL’s strong predictive ability, capable of not only forecasting drug-target interactions and binding affinities but also pinpointing specific binding regions.

### Performance of DrugDL on the molecular properties and toxicities prediction tasks

We first evaluated the performance of DrugDL using five types of benchmark datasets of physicochemical properties of drug molecules: BACE, BBBP, Estimated Solubility (ESOL), Lipophilicity, and Free Solvation (FreeSolv). We compared DrugDL with three state-of-the-art methods for predicting the properties of drug molecules: MoleculeNet^53^, HiMol^54^, and HimGNN^55^. The prediction results of each model are shown in Fig. 3a. Overall, the DrugDL model demonstrated better performance than other baseline models in all evaluation metrics. On the BACE dataset, the AUC, ACC, Recall, and Specificity of DrugDL were 0.9237, 0.9249, 0.9034, and 0.9465 respectively, all higher than those of other models. Similarly, on the BBBP dataset, DrugDL also reached the highest values for these four indicators. Although the prediction effects of HiMol and MoleculeNet were similar, and HimGNN was slightly better than the former two in terms of AUC and ACC, their performances were still inferior to that of DrugDL. On the ESOL, Lipophilicity, and FreeSolv datasets for regression tasks, DrugDL also performed outstandingly.

**Fig. 3.**
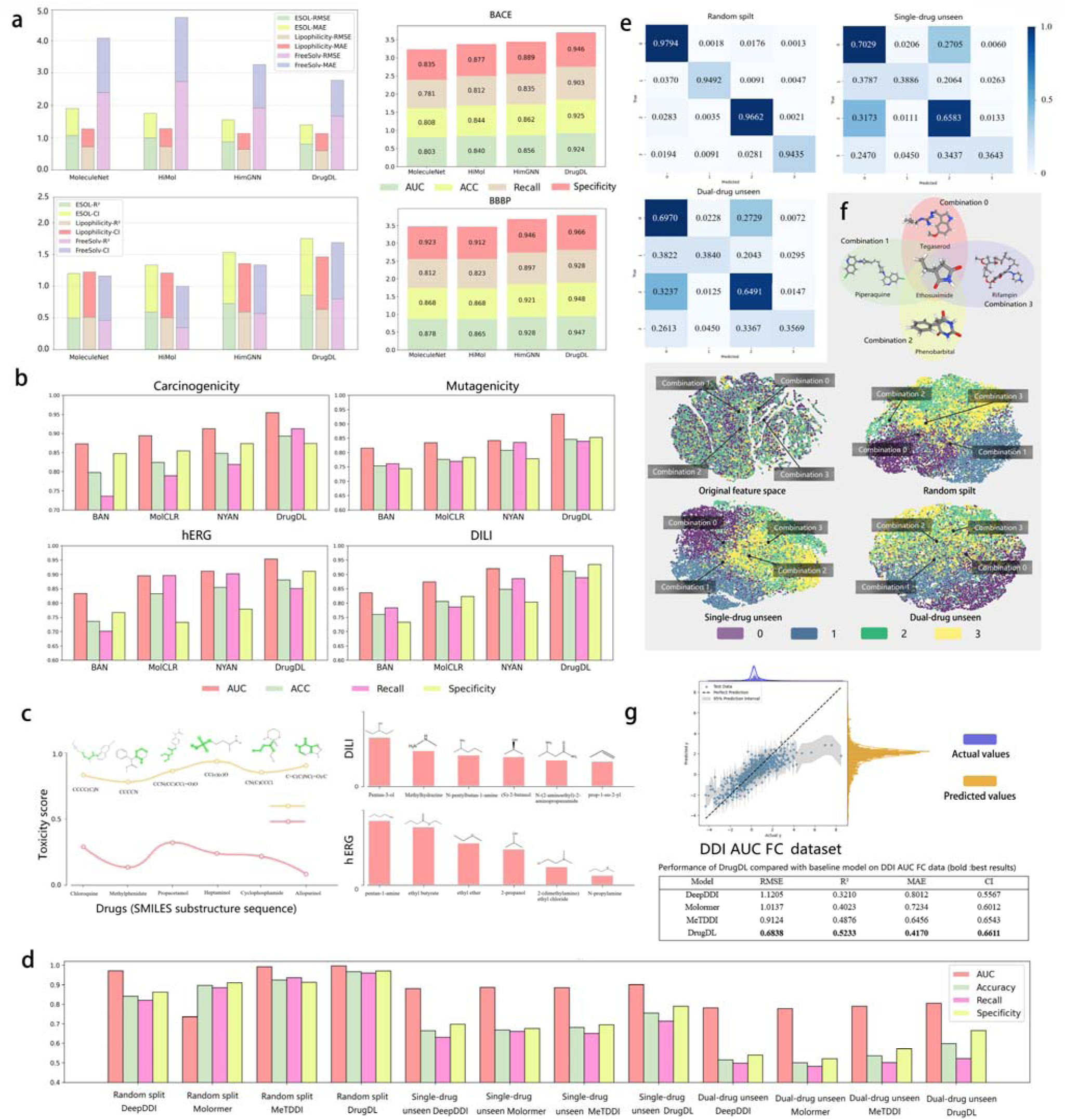
Performance evaluation on the drug safety prediction task. **a**, Evaluation of the performance of DrugDL and baseline models on datasets of physicochemical properties of drug molecules (BACE, BBBP, ESOL, Lipophilicity, and FreeSolv). **b**, Evaluation of the performance of DrugDL and baseline models in terms of AUC, ACC, Recall and Specificity on datasets of toxicity of drug molecules (carcinogenicity, mutagenicity, DILI and hERG). **c**, Comparison of toxicity before and after the removal of substructures identified by DrugDL and analysis of sensitive areas. **d**, Evaluation of the performance of DrugDL and each baseline model in terms of AUC, ACC, Recall and Specificity for the DDI dataset in three scenarios: randomly split dataset, single-drug unseen dataset and dual-drug unseen dataset. **e**, Classification confusion matrices of DrugDL in three DDI dataset segmentation scenarios. **f**, Taking the drugs Ethosuximide, Tegaserod, Piperaquine, Phenobarbital and Rifampin as examples, the prediction visualization results of DrugDL in three DDI dataset segmentation scenarios. **g**, Performance comparison between DrugDL and baseline models based on DDI AUC FC value dataset and prediction visualization results.

In drug toxicity prediction, we compared our DrugDL model with three advanced baseline models—BAN^56^, MolCLR^47^, and NYAN^57^—across four key toxicity datasets: carcinogenicity, mutagenicity, human ether-à-go-go gene (hERG) cardiotoxicity, and drug-induced liver injury (DILI). As shown in Fig. 3b, DrugDL significantly outperformed all baseline models in AUC, ACC, and Specificity metrics, demonstrating its strong predictive ability and stability. This highlights DrugDL’s advantage in capturing drug molecule toxicity features. On the DILI dataset, while DrugDL’s Recall was slightly lower than NYAN and MolCLR, these baseline models achieved higher Recall by classifying most data as positive, which increased false positives and reduced Specificity and overall ACC. In contrast, DrugDL effectively balanced Recall and Specificity, reducing false positives and enhancing model reliability and practicality. Thus, DrugDL offers greater clinical value in comprehensive performance, particularly in minimizing unnecessary diagnoses and misjudgments. Additionally, using DrugDL’s substructure extraction method, we identified substructure sequences crucial for toxicity judgment and presented them in Supplementary Table 1, including substructure images, toxicity, and SMILES representations. When we removed these substructures and conducted toxicity tests, the results (Fig. 3c) confirmed that the model’s identified substructures reduced compound toxicity. Fig. 3c also highlights the six substructures most sensitive to different toxicities, providing valuable insights into drug molecule toxicity mechanisms.

### Performance of DrugDL on the DDI and pharmacokinetic impact prediction tasks

In the DDI prediction task, we conducted an in-depth comparison between DrugDL and three baseline models, DeepDDI^58^, Molormer^59^, and MeTDDI^48^. The performance results of DrugDL and the baseline models on the DDI benchmark dataset are shown in Fig. 3d. Firstly, in the randomly split dataset, the DrugDL model demonstrated excellent performance. In all evaluation metrics, DrugDL significantly outperformed all baseline models. However, when we divided the dataset into single-drug unseen and dual-drug unseen subsets, the prediction performance of all models decreased significantly. Nevertheless, the DrugDL model still maintained excellent prediction performance on these two types of datasets. The prediction classification results on the three classification datasets are shown in Fig. 3e, and the classification results presented by its confusion matrix correspond to those in Fig. 3d. Fig. 3f further shows the t-SNE visualization results of the DrugDL model in the three classification tasks and the original feature space. We centered on the drug Ethosuximide to represent its four relationships with other drugs (Tegaserod, Piperaquine, Phenobarbital, and Rifampin). By observing the t-SNE visualization results, we can find that the DrugDL model successfully separated the four types of data in the randomly split dataset, and the boundaries between various types of data were relatively clear. At the same time, we also paid attention to the t-SNE visualization results of the unseen datasets. Although the model’s prediction results on this part of the data were not as good as those on the randomly split data, these data points still showed a similar clustering trend to some extent. This further demonstrates the advantage of the DrugDL model in dealing with the cold start problem.

In addition, to comprehensively evaluate the performance of various models in DDI prediction, we also took into consideration the area under the plasma time concentration curve fold change (AUC FC) values, which are used to measure the pharmacokinetic impact. As shown in Fig. 3g, the prediction performance of the DrugDL model on the DDI AUC FC value data was also better than that of the baseline models. By observing the scatter plot, we can clearly see that the distribution of the predicted values is roughly the same as that of the actual values, and the 2D points composed of the actual values and the predicted values are basically distributed on the diagonal. This result not only proves the effectiveness and accuracy of the DrugDL model but also provides strong support for its application in fields such as drug R&D and drug interaction evaluation.

### DrugDL identifies potential inhibitors for Anti-SARS-CoV-2 and metabolic enzymes

To deeply explore the practical effectiveness of the DrugDL model in real world drug development and clinical applications, we systematically evaluated the performance of DrugDL and other baseline models (MolCLR^47^, NYAN^57^, and ImageMol^21^) in two key areas: anti-SARS-CoV-2 drug development and metabolic enzyme inhibitor identification, verifying its potential in real world application scenarios.

Firstly, in the field of anti-SARS-CoV-2 drug development, SARS-CoV-2 is the source of the Coronavirus Disease 2019 pandemic. In the life cycle of SARS-CoV-2, the 3CL protease is a crucial part of virus replication^60,61^. We evaluated the antiviral potential of drugs by predicting and analyzing whether drug molecules can bind to the 3CL protease and inhibit its activity. The relevant results show (Fig. 4a) that in the AUC and AUPR, DrugDL demonstrated significant advantages over other baseline models, achieving scores of 93.24 and 93.23 respectively. In addition, the features of drug molecules extracted by DrugDL achieved good clustering of whether the molecules belong to anti-SARS-CoV-2 drugs.

**Fig. 4.**
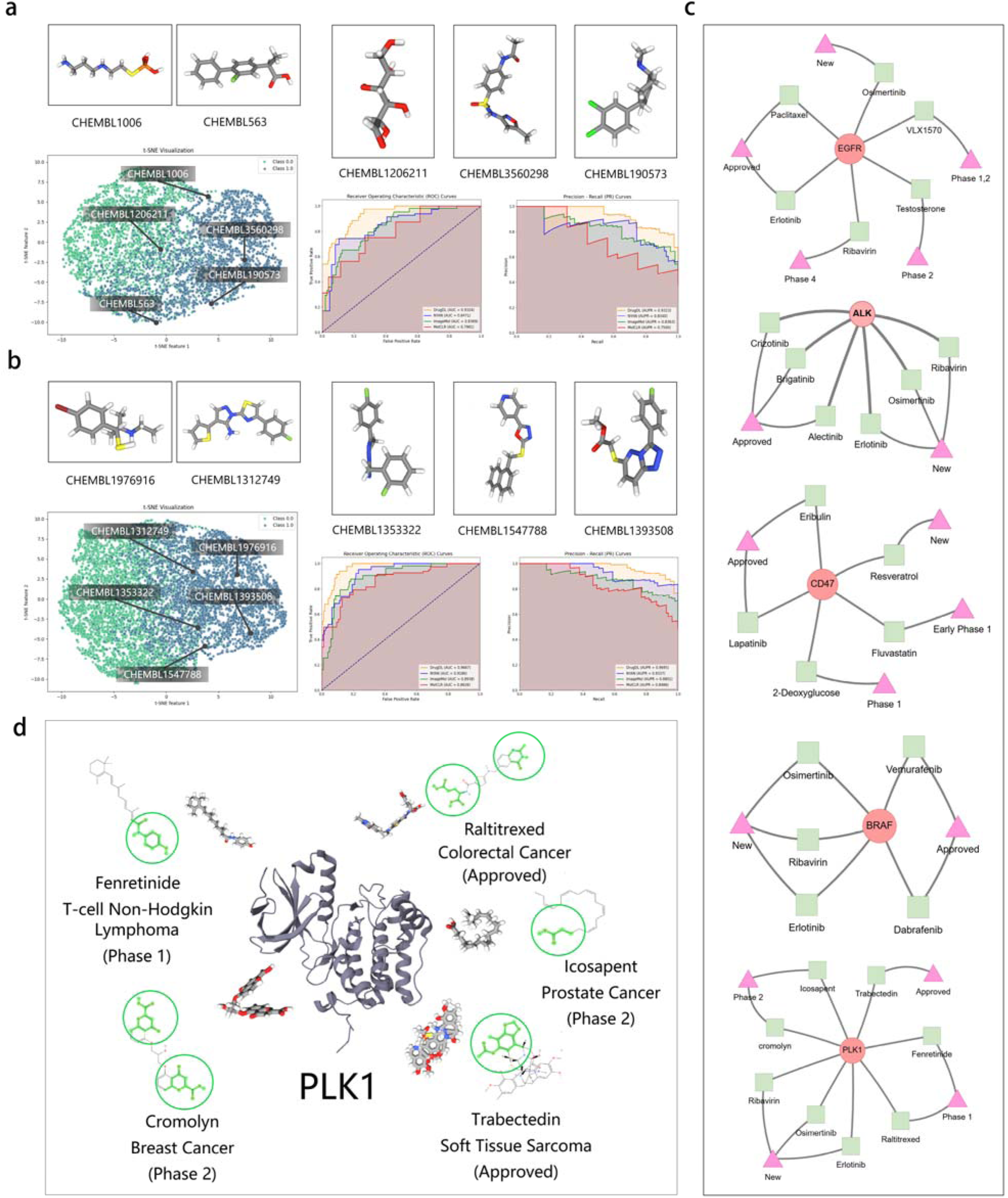
Evaluation of the practical efficacy in real world drug R&D and clinical applications. **a**, Evaluate the performance of DrugDL and baseline models on the anti-SARS-CoV-2 inhibitor dataset. **b**, Evaluate the performance of DrugDL and baseline models on the metabolic enzymes (CYP2C9) inhibitor dataset. **c**, Analysis of DrugDL in the prediction of potential drugs for tumor targets (BRAF, ALK, EGFR, CD47, and PLK1). **d**, Visualization results of drug molecules interacting with the PLK1 target identified by DrugDL.

Next, in the field of metabolic enzyme inhibitor identification, the CYP2C9 metabolic enzyme, as a representative metabolic enzyme, plays an important role in the metabolism of many clinically commonly used drugs^62,63^. Through the prediction and analysis of whether a drug is a CYP2C9 inhibitor, DrugDL also demonstrated its excellent performance. In the ROC curve evaluation (Fig. 4b), the AUC value of DrugDL reached 0.9687, significantly higher than that of other baseline models. In the PR curve evaluation, the AUPR value of DrugDL was also 0.9695, which means that DrugDL can still effectively maintain a balance between high Precision and Recall even in the case of unbalanced samples. In conclusion, DrugDL has demonstrated its cross-field application potential in the two different fields of anti-SARS-CoV-2 drug development and CYP2C9 enzyme inhibitor identification.

### DrugDL predictive identification of cancer therapeutics and mechanistic analysis

To further evaluate the reliability of the DrugDL model in practical application scenarios, we selected multiple tumor-related targets for in-depth case studies. These targets include BRAF (B-Raf proto-oncogene)^64^, ALK (anaplastic lymphoma kinase) ^65^, EGFR (epidermal growth factor receptor) ^66^, CD47 (a key transmembrane protein) ^67^, and PLK1 (an important protein in cell cycle regulation) ^68^. For each target, we first pre-trained DrugDL using a benchmark dataset, and then predicted the drugs that might interact with these targets on the dataset.

As shown in Fig. 4c, we presented several drugs with high predicted interaction probabilities for each target. In the list of drugs predicted to interact with the BRAF target, two drugs, Vemurafenib and Dabrafenib, have been approved for the treatment of melanoma and other related cancers, which initially validates the effectiveness of our prediction method^69,70^. Although there is currently no direct clinical trial evidence, our prediction results suggest that Erlotinib and Osimertinib may have significant interactions with the BRAF target. It is worth mentioning that these two drugs have shown potential anti-cancer activities in multiple studies and have been supported by clinical trials. These findings indicate that the new drugs predicted by DrugDL to interact with specific targets are promising candidates for further exploration of cancer treatment. In the prediction for the PLK1 target, Fenretinide is in the Phase 1 clinical trial for T-cell non-Hodgkin lymphoma^71^, and Cromolyn and Icosapent are in the Phase 2 clinical trials for breast cancer and prostate cancer^72,73^. For the CD47 target, the predicted drug Fluvastain has entered the early Phase 1 clinical trial for malignant melanoma^74^.

In addition, we conducted an analysis of the DTI mechanisms for five drug molecules (Fenretinide, Cromolyn, Trabectedin, Icosapent, and Raltitrexed) that bind to the PLK1 target. Fig. 4d depicts the 3D structures of these molecules, as well as the attention weights of the corresponding atoms and motifs. We highlighted the substructures with large attention weights in green. These highlighted parts help us deeply analyze and understand the complex and subtle interaction examples between drugs and targets, and enhance our understanding of the key functional groups in drug molecules.

### Experimental validation of EGFR and ALK-targeting drug candidates and toxicity assessment

To assess virtual drug screening, this study evaluated DrugDL-identified candidates for EGFR and ALK targets. Fig. 5a shows the top 10 predicted drugs for each wild-type protein. DrugDL deemed 7 of 10 EGFR and 5 of 10 ALK inhibitors safe. Among safe EGFR inhibitors, Erlotinib, VLX15703, Testosterone, and Paclitaxel were validated externally. PF-04217903, BMS-345541, and TAK-285, novel safe drugs, bound both EGFR and ALK safely. Further validation was conducted using molecular docking experiments (AutoDock Vina).

**Fig. 5.**
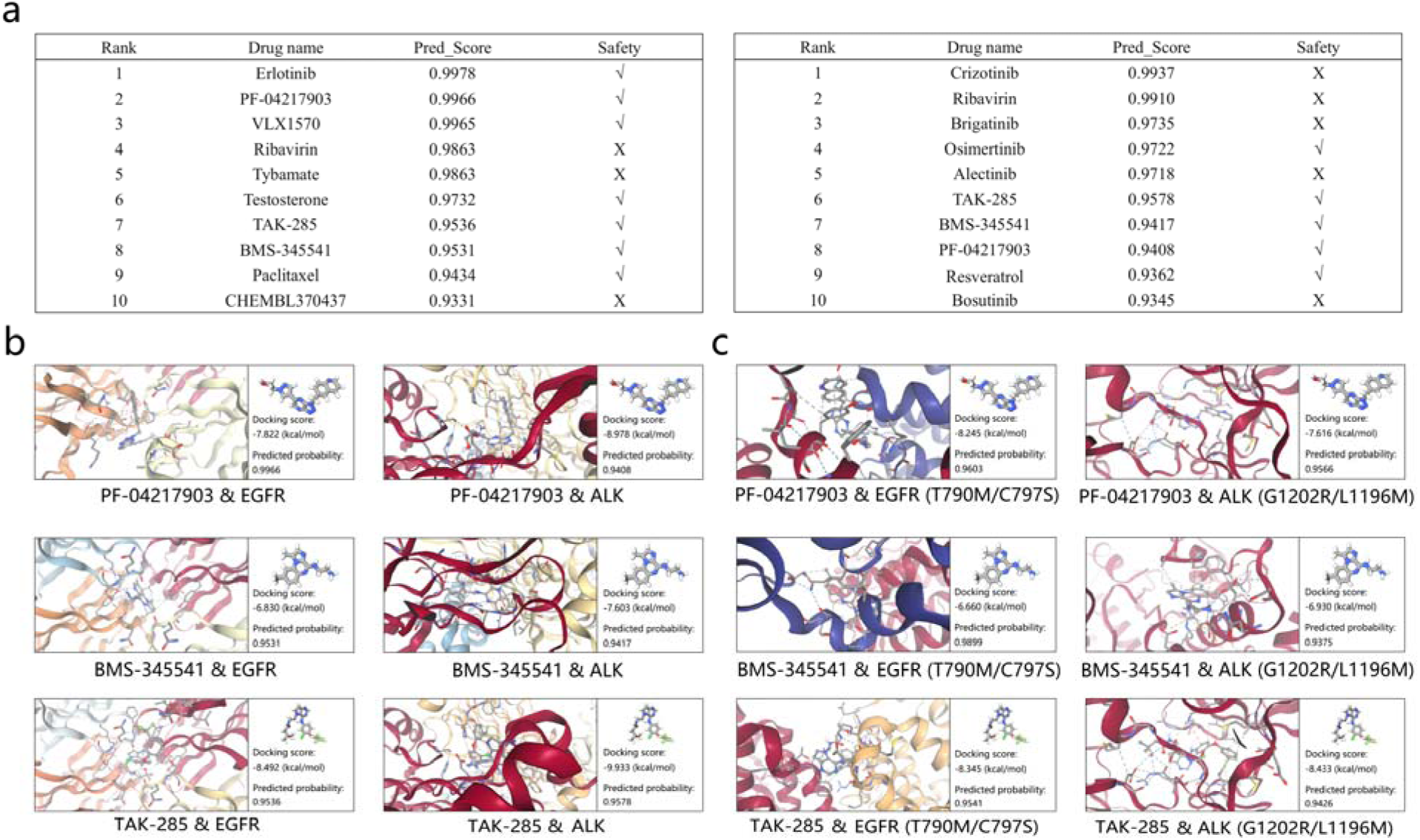
Experimental verification and toxicity evaluation results of EGFR and ALK targeted drug candidates identified by DrugDL. **a**, Results of DrugDL predicting top 10 potential drugs binding to EGFR targets and top 10 potential drugs binding to ALK targets. **b**, The docking poses and scores of the predicted interactions between three potential dual-target drugs (PF-04217903, BMS-345541, and TAK-285) and the EGFR and ALK targets.

Molecular docking analysis (Fig. 5b) revealed that the binding free energies (ΔG) of the three candidate drugs with wild-type EGFR and ALK proteins ranged from −6.8 to −10.0 kcal/mol. According to thermodynamic criteria (ΔG < 0), these values indicate that the binding processes between the drug molecules and target proteins are spontaneous and exhibit moderate to strong affinity. Notably, PF-04217903 and TAK-285 showed particularly strong ALK affinity (ΔG < −8.9 kcal/mol), while TAK-285 also bound well to EGFR (ΔG = −8.492 kcal/mol), further highlighting the relative advantages of these drugs in binding to specific targets.

However, in clinical practice, tumor cells often undergo genetic mutations, leading to alterations in the structure and function of target proteins. In recent years, EGFR mutations such as T790M and C797S have emerged as resistance-causing mutations for third-generation drugs. The T790M mutation alters the spatial structure of the ATP-binding pocket in EGFR, impeding drug-target binding, while the C797S mutation may disrupt covalent binding between drugs and EGFR, rendering the drugs ineffective. Similarly, ALK mutations like G1202R and L1196M can alter the structure of ALK, making it difficult for drugs to bind to the mutated ALK protein and reducing their antitumor activity. Consequently, there is an urgent need to identify novel drug candidates capable of addressing target mutations.

Thus, we predicted the interaction scores of the three dual-target drugs with two mutated targets using DrugDL. The results indicated that all three drugs interacted with both target proteins. Further validation was conducted through docking experiments, which revealed that ΔG of the three candidate drugs with the double-mutated EGFR and ALK proteins ranged from −6.6 to −8.5 kcal/mol, suggesting strong affinity between the drug molecules and target proteins. Additionally, the results of molecular docking experiments between the candidate drugs and other mutated EGFR and ALK proteins are presented in the Supplementary Table 3.

## Discussion

We introduce DrugDL, a cross-modal deep learning framework for drug molecule representation and multi-property prediction. It integrates multi-level drug and target protein data using multi-channel GNNs and CNNs, analyzing multi-scale interactions via cross-modal learning and single-modal enhancement while preserving feature attributes to prevent information loss. Our experiments show DrugDL outperforms existing methods in predicting DTIs, DTAs, DTBRs, drug physicochemical properties, toxicity, and DDIs. Moreover, it identifies crucial molecular substructures with pharmacological insights, aiding drug researchers in screening key action sites and uncovering local interaction mechanisms. DrugDL’s practical efficacy is demonstrated in real-world R&D and clinical applications, providing theoretical support for drugs at various clinical trial stages by predicting cancer-target interactions and experimentally validating EGFR/ALK-targeted candidates’ efficacy and safety.

We observe that while DrugDL’s core paradigm involves representing drug molecule features and predicting downstream molecular properties by analyzing DTIs, the availability of such interaction data is not the primary bottleneck limiting DrugDL’s efficacy. Specifically, the single-modal feature enhancement module in DrugDL can independently model drug and target modalities, fully preserving their distinct features and preventing information loss during multi-modal fusion. Crucially, this module enables indirect inference of key information through intra-modal information transfer and cross-modal collaborative learning, even with limited direct DTI data, by explicitly capturing correlations within modal data (e.g., drug-drug and target-target similarities). Compared to prior work, DrugDL is less constrained by data availability, a advantage fully validated by our model evaluations in cold and warm start scenarios.

In deep learning-based biological discovery, where black-box prediction mechanisms are unclear, DrugDL excels in parsing pharmacologically informative substructures. It converts drug molecules into graph representations using a pharmacophore-motif-guided substructure splitting approach, extracting multi-scale pharmacology-related substructures (e.g., pharmacophores, toxicophores). By leveraging attention mechanisms and interpretability techniques, DrugDL quantifies each substructure’s contribution to pharmacological indicators and generates hypotheses with mechanistic insights. Substructure analysis serves as an “interpretability anchor” for deep learning models, partially elucidating their prediction logic and guiding virtual screening and molecular design optimization to expedite drug discovery. DrugDL’s success in EGFR/ALK-targeted drug discovery validates its feature extraction and decision-making model. However, despite its interpretability potential in drug property prediction, most model outputs require wet-lab experimental validation. Future research will employ advanced technologies to systematically verify the biological credibility of DrugDL-identified substructures, aiming to enhance drug structure optimization and achieve simultaneous improvements in drug safety and efficacy.

## Methods

### The workflow of DrugDL

DrugDL consists of three main components: (1) Cross-modal interaction learning that accurately captures the potential and complex relationships between drugs and targets; (2) Single-modal feature enhancement used to strengthen the characteristic attributes of drugs and targets themselves and prevent information loss; (3) Downstream drug molecule property prediction tasks via machine learning and deep learning.

### Cross-modal Interaction Learning Module

For each drug molecule, we construct two types of graphs: one is a drug atom graph with atoms as nodes and chemical bonds as edges, and the other is a drug motif graph with motifs as nodes and edges constructed between nodes based on multiple principles. These two graphs reveal the structural features and interaction relationships of drug molecules from different perspectives. The atom graph *G_a_* = (*V_a_*, *E_a_*), where *V_a_* represents the set of atom nodes and *E_a_* represents the set of edges composed of chemical bonds between atoms. A drug molecule contains n atoms, and each atom *i* (*i=1,2,…,n*) corresponds to a node *v_i_* ε *V_a_* in the graph. Each node in the atom graph will precisely reflect the local structural features of the atom. The motif graph *G_m_* = (*V_m_*, *E_m_*), where *V_m_* represents the set of nodes composed of motifs and *E_m_* represents the set of edges between nodes^75^.

To extract deeper information in drug molecules, we use multi-channel GNNs to learn the representation of drugs. Specifically, in addition to using two parallel GATs to extract features from the drug atom graph and the motif graph, we also use a GCN with shared weights to extract the consistent features of the two graphs.

In the process of target feature representation learning, to comprehensively and accurately extract the adjacent information in the protein sequence, we choose to use a CNN for protein sequence feature extraction. First, we perform a k-mer encoding operation on the amino acid sequence of the protein. This encoding method lays a solid foundation for subsequent feature extraction, enabling the sequence information to be processed by the network in a more suitable form. Subsequently, we carefully design a three-layer CNN block specifically for protein feature extraction. In each convolutional layer, we deploy multiple convolutional kernels. These convolutional kernels, like sensitive detectors, each focus on learning the embedding representation of a specific region of the sequence. For each protein sequence, the convolutional kernels perform convolutional calculations in an orderly manner. Each convolutional kernel is tasked with extracting the information of a specific segment in the sequence, thus mining the potential features of the sequence from different perspectives. Inspired by the residual network, we ingeniously connect a feature aggregation module between different convolutional layers. This module is composed of a pooling layer and an activation function. We set the pooling layer as a non-linear pooling function, specifically, the “max-pooling” operation. Under the max-pooling operation, the length of the sequence feature vector is halved after each pooling, which effectively reduces data redundancy. The convolutional layer connected after the pooling layer is equivalent to performing a linear weighting on the result of the non-linear function. This design not only strengthens the function of the pooling layer in reducing information redundancy but also minimizes the information loss caused by the pooling operation. Finally, we splice the result of the pooling operation with the result of the convolutional operation and successfully obtain the high-level embedding features of the protein.

Next, we will use the contrastive learning loss to align the features of the two modalities of drugs and targets. We assume that there are *N* pairs of drug-target data, and their feature representations are (*D_i_*, *T_i_*). When there is an interaction between drug *i* and target *i*, we regard it as a positive sample; otherwise, it is a negative sample.

### Single-modal Feature Enhancement Module

In the single-modal feature enhancement module, we adopt unique similarity calculation methods and loss functions to preserve the internal relationships existing in each modality. For the drug feature matrix and the target matrix after feature extraction, we use matrix SVD to extract the principal components in the feature matrix, effectively eliminating redundant information while accurately retaining key information. Then we introduce the Gaussian kernel function to measure and calculate the similarities between different drugs and between different targets in the embedded feature space. At the same time, in the original feature spaces of drugs and targets, after normalization, we also use SVD for information extraction and the Gaussian kernel function to calculate the internal similarities of the data in each modality. For the similarities of each modality’s data in the original feature space and the embedded feature space, we measure them using the consistency loss of the similarity matrix, and sum the loss values of the two modalities’ data to obtain the final single-modal preservation loss.

### Downstream Drug Molecule Property Prediction

In the crucial downstream tasks of drug discovery, we have constructed a multi-task prediction framework that covers core scenarios such as DTI, DTA and DTBR prediction, prediction of drug physicochemical properties, toxicity assessment, and DDI. For tasks that require cross-modal integration (such as DTI, DTA, and DTBR), we fuse the representations of drugs and targets into a joint feature vector through feature splicing and input it into the prediction model. For single-modal tasks (such as prediction of physicochemical properties and toxicity), we directly use the drug molecule features as input. To achieve efficient modeling, we adopt a hybrid method framework. For structured features, in addition to training traditional machine-learning models such as Support Vector Machines (SVM), Random Forest, and Logistic Regression, we also design fully-connected neural networks (FCNN) with different architectures for in-depth feature mapping. The FCNN we designed contains multiple hidden layers, uses the ReLU activation function and Batch Normalization to accelerate convergence, and finally generates prediction probabilities through a Sigmoid or Softmax output layer.

### Benchmark Datasets

To comprehensively evaluate the model performance, we have integrated multiple authoritative public datasets to construct a multi-task benchmark system, covering core tasks such as DTI, DTA and DTBR prediction, prediction of physicochemical properties, toxicity assessment, and prediction of DDI. For the DTI task, we use the BindingDB^76^, BioSNAP^77^, and Human datasets^78^. Among them, BindingDB contains more than 50,000 verified interaction data of small molecules and proteins. BioSNAP is constructed based on DrugBank and contains information on nearly 30,000 drug molecule-protein interactions. The Human dataset focuses on the prediction of human-specific interactions. For binding affinity prediction, we use the Davis^79^ and KIBA^80^ benchmark datasets. Davis provides dissociation constant data of the kinase family, and KIBA integrates multi-source biological activity indicators and standardizes the scores. For binding site prediction, we use the sc-PDB dataset, which contains 16,000 experimentally verified protein-ligand binding sites**^Error! Reference source not found.^**.

For the prediction of physicochemical properties, we have integrated five subsets of MoleculeNet^53^: human β-secretase inhibitor (BACE), blood-brain barrier permeability (BBBP), Estimated Solubility (ESOL), Lipophilicity, and Free Solvation (FreeSolv), covering nearly 10,000 drug molecules in total. Toxicity assessment includes four types of tasks: carcinogenicity, mutagenicity, human ether-à-go-go gene (hERG) cardiotoxicity, and drug-induced liver injury (DILI). The carcinogenicity and mutagenicity data integrates the CPDB^82^, CCRIS^83^, and ISSCAN^84^ databases, covering nearly 10,000 molecules. The hERG toxicity dataset is constructed based on ChEMBL^85^, PubChem^86^, and BindingDB^76^, containing 20,000 molecules and dividing toxicity labels according to whether the IC50 value is lower than 10 μM. The liver injury data integrates the LTKB, LiverTox, and Hepatox databases^87^, covering the clinical adverse reaction records of nearly 10,000 drugs.

For the study of DDIs, we use DrugBank 5.1.8^88^ to construct a four-class classification task and a regression task. The four-class classification task defines two-way labels for drug metabolism inhibition and enhancement relationships, and the regression task quantifies the pharmacokinetic effects through the area under the plasma time concentration curve fold change (AUC FC) values. All datasets have been processed through deduplication, arbitration of conflicting labels, and prioritization of experimental verification to ensure data quality and reproducibility.

### Baselines

To systematically evaluate the performance of the DrugDL framework, this study constructs a multi-level baseline system from the perspective of task adaptability. At the level of drug molecule representation, traditional molecular fingerprint methods are compared: Morgan fingerprints (based on topological encoding of molecular substructures), ECFP (Extended Connectivity Fingerprints), PubChem fingerprints (functional group descriptors), MACCS fingerprints (predefined key substructures), and Pharmacophore ErG fingerprints (spatial feature encoding of pharmacophores), to verify the distinctiveness and information density of the features generated by DrugDL.

For the prediction task of DTI, three types of cutting-edge models are selected: DrugBAN^30^ captures the local interaction patterns between drugs and targets through a bilinear attention network; ZeroBind^31^ achieves cold start prediction based on the prior knowledge of protein structures; and PSICHIC^32^ integrates multi-source biological networks to construct a heterogeneous graph attention model. For the prediction of DTA, MFR-DTA^42^ (multi-scale feature recombination for optimizing binding free energy modeling), KDBNet^39^ (a two-tower architecture enhanced by knowledge distillation), and MMD-DTA^40^ (multi-modal disentangled representation learning) are compared. For the DTBR prediction task, TransformerCPI^52^ (self-attention-driven residue localization), MFR-DTA (extending affinity prediction to spatial site recognition), and MMD-DTA (a cross-modal feature alignment framework) are introduced to verify the representation ability of DrugDL for the microscopic binding mechanism.

For the prediction of drug physicochemical properties, MoleculeNet^53^ (a standardized multi-task learning framework), HiMol^54^ (a hierarchical GNN), and HimGNN^55^ (a heterogeneous information fusion model) are used as baselines to evaluate the generalization ability of DrugDL features in tasks such as solubility and lipophilicity. In the toxicity prediction task, BAN^56^ (bidirectional attention modeling of toxic fragments), MolCLR^47^ (contrastive learning to enhance molecular representation), and NYAN^57^ (a network for inferring toxic metabolic pathways) are compared, covering four scenarios: carcinogenicity, mutagenicity, hERG cardiotoxicity, and liver injury. For the prediction of DDI, DeepDDI^58^ (substructure interaction modeling), Molormer^59^ (a Transformer-based molecular pair encoder), and MeTDDI^48^ (a multi-view pharmacokinetic impact prediction framework) are selected. The robustness of the model is verified by predicting the relationships of drug metabolism inhibition and enhancement and the AUC FC value. All baseline models strictly reproduce the experimental settings in the original papers to ensure the fairness of the comparison.

### Experimental settings

DrugDL and the above-mentioned baseline methods will be evaluated under specific sample processing strategies and different validation settings. To address the issue of sample imbalance, we have implemented targeted treatment measures. We oversample the positive samples and undersample the negative samples simultaneously, ensuring that the number of positive samples reaches a consistent level after oversampling, so as to balance the distribution of positive and negative samples and provide a more balanced data foundation for subsequent model training. Subsequently, we divided the dataset according to strict standards and carried out experiments based on this. Specifically, we extracted 80% of the positive and negative sample pairs from the positive and negative sample sets as the training set, and reserved the remaining 20% of the sample pairs as the test set to evaluate the prediction performance of the model.

When dividing the DTI dataset, we adopted two methods: random splitting and cold start splitting^30^. Random splitting means that the dataset is randomly divided into a training set and a test set according to a preset ratio; while cold start splitting ensures that the drug set and the protein set in the test set do not appear in the training set while maintaining the splitting ratio, so as to avoid the model from relying too much on known features. This approach enables the model to demonstrate a more realistic prediction ability when facing test data, rather than just making inferences based on the learned features of drugs and proteins.

In the research field of DDIs, we followed the data preprocessing standards of MeTDDI and reversed the order of the drug pairs with data labels 1 and 2, thus obtaining the information of drug pairs with labels 3 and 4. Specifically, label 1 represents that when drug *d*_1_ and drug *d*_2_ are used in combination, it can reduce the metabolism of drug *d*_1_, and label 2 represents an increase; labels 3 and 4 represent the reduction and increase of the metabolism of drug *d*_2_ when drug *d*_1_ and drug *d*_2_ are used in combination, respectively. When splitting the DDI dataset, in addition to the random splitting method, we also introduced more detailed cold start splitting strategies of single-drug unseen and dual-drug unseen, enabling the model to make predictions on individual drugs and drug pairs that it has never seen before, helping us to more comprehensively evaluate the generalization ability of the model.

### Evaluation metrics

In this study, we comprehensively evaluate the performance of the model in classification and regression tasks with the help of multiple evaluation metrics. In terms of classification tasks (such as DTI prediction, prediction of some drug physicochemical properties, toxicity prediction, etc.), the test results are presented and analyzed under four core evaluation metrics, namely Accuracy (ACC), area under the receiver operating characteristics curve (AUC), Specificity, and Recall. For multi-classification scenarios (DDIs), in order to provide a more detailed performance analysis, we further introduce the average Specificity and average Recall of each category. In regression tasks (such as DTA prediction, drug AUC FC values, and prediction of some drug physicochemical properties), we select Root Mean Square Error (RMSE), Coefficient of Determination (R²), Mean Absolute Error (MAE), and Consistency Index (CI) as key evaluation metrics. In addition, in some tasks (DTA prediction), to ensure a fair comparison with the baseline model, we use the variant 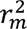 of the squared correlation coefficient value for evaluation and analysis.

## Ethics declarations

### Ethical approval

As publicly accessible documents have been used for the current study, ethical approval is not required for this study.

